# Hidden Diversity in Yeast tRNAs: Comparative Genomics and Modification Mapping in a Eukaryotic Subphylum

**DOI:** 10.64898/2026.03.20.712421

**Authors:** Lauren Dineen, David Wilson, Abigail Leavitt LaBella

**Affiliations:** Department of Bioinformatics and Genomics, University of North Carolina at Charlotte, North Carolina Research Campus, Kannapolis, NC 28081; Department of Infectious Disease, Imperial College London, London SW7 2AZ, United Kingdom; Department of Bioinformatics and Genomics, University of North Carolina at Charlotte, Charlotte, NC 28223; Center for Computational Intelligence to Predict Health and Environmental Risks, University of North Carolina at Charlotte, Charlotte, NC 28223

**Keywords:** tRNA, tRNA modification, yeast, genomics

## Abstract

tRNA are adapter molecules with an integral role in translation and further roles in stress adaptation. Processing of tRNA is tightly regulated and includes the enzymatic addition of several post-transcriptional modifications that are required for translation efficiency, recognition, selective translation, and structure. We currently lack a multi-species wide view of tRNA modifying enzymes across eukaryotes. Here, we performed a comparative analysis of tRNA gene sequence, modification enzymes, and modification profiles across the Saccharomycotina subphylum. We employed machine learning methods to explore tRNA sequence conservation and to annotate modifying enzymes known to exist in fungi, humans, and prokaryotes. We then applied Nano-tRNAseq to three species (*Saccharomyces cerevisiae, Hanseniaspora uvarum, and Yarrowia lipolytica*) to profile modification signatures and compare modification patterns. We identified substantial lineage-specific conservation of tRNA sequences despite the highly conserved tRNA structure. We found significant variation in tRNA modifying enzyme repertoires across Saccharomycotina, including lineage-specific losses, and annotated a prokaryotic-associated enzyme, *tilS*. Integrating genomic and sequencing data enabled us to link enzyme repertoires with tRNA gene sequences. tRNA sequencing revealed distinct modification signatures across the three focal species, and further analysis using General Linearized modelling suggested tRNA enzyme loss is associated with target tRNA nucleotide absence in gene sequences. This work provides the first integrated view of tRNA gene and modification diversity in eukaryotes and expands the field of tRNA diversity in fungi.

## Introduction

Transfer RNAs (tRNA) are found in all known organisms and play a critical role in the essential process of translation in the Central Dogma of biology. The primary role of tRNA is to deliver amino acids cognate to the messenger RNA codon to the ribosome during translation. Despite the integral role of tRNA, there remains a remarkable knowledge gap in fundamental tRNA biology due to the previously accepted notion that tRNAs were passively expressed background molecules. In recent decades, however, advances in sequencing technology and genomics have opened the field of tRNA biology and challenged these assumptions to reveal that tRNAs are important regulatory molecules that influence translation dynamics and other cellular processes ^1–7^.

tRNA molecules are ancient—their function in translation, and secondary ‘cloverleaf’ and tertiary ‘elbow’ structure are conserved across the tree of life. Despite their small size, ∼70-90nt, tRNA sequences contain a wealth of information such as internal promoter regions, introns, and aminoacylation and modifying enzyme identifiers. tRNAs are classified by amino acid group, anticodon, and unique gene sequence; these are termed isodecoders. tRNA can exist in the genome as exact duplications, or more than one isodecoder with unique gene sequences can exist within a group. Any given genome contains variable levels of tRNA at each classification, with some genomes containing the minimum viable set of tRNAs ^8^ and others containing hundreds of isodecoders ^9,10^. The full functional significance of this diversity remains unclear. Previous work has shown that tRNA gene copy number tends to correlate with genomic codon usage ^11–14^. This suggests that regulation of the tRNA pool, along with non-canonical functions, may influence the diversity of isodecoders between species ^9,15–17^.

tRNAs are highly regulated and require complex pre-processing steps to reach maturity. The tRNA pool, referring to the pool of available tRNA in a cell, is dynamic in nature and is regulated by differential expression, post-transcriptional processing, and degradation/decay pathways ^17–21^, although further novel mechanisms are likely to exist. Further regulated pre-processing steps involve splicing, addition of the 5’ CCA, post-transcriptional modification and aminoacylation ^22^. All of which create a complex and multi-layered system of regulation with direct effects on protein synthesis.

A key part of translation is the presence of wobble codons in gene transcripts, so called because there is not a cognate tRNA encoded in the genome. To translate these codons, post-transcriptional modifications to the tRNA anticodon expand the decoding ability of a tRNA to decode more than one codon. Examples of stress-induced translational regulation via modifications at the wobble base have been described ^3,23^. In these studies, the addition of wobble-base pairing modifications drives increased translation of wobble codon enriched transcripts to aid cellular stress response. tRNA are highly modified across the total structure, with an average of 13 modifications per molecule described in eukaryotes ^24^. tRNA modifying enzymes (tME) are position and nucleotide-specific, and require diverse substrates and multi-enzymatic pathways ^22,25^. Furthermore, the presence of certain modifications can determine the addition of others ^26–28^. Eukaryotic tMEs are known to ensure tertiary structure, maintain translation, regulate tRNA fragment biogenesis, and enable identification by amino-acylation enzymes ^29–35^. These results derive mainly from model organisms, and the full spectrum of eukaryotic tMEs has yet to be uncovered.

Despite leaps in the field, there remains a significant gap in knowledge of the fundamental tRNA biology in eukaryotes. This includes the mechanisms behind tRNA pool dynamics and regulation, complete roles of tRNA modifications and modifying enzymes and non-canonical roles of tRNA. Secondly, limitations in tRNA sequencing technology result from short length fragments, tertiary folding, sequence similarities, and their heavily modified nature ^36,37^. At the forefront of this gap is the absence of substantially powered comparative studies between species; as a result, the current understanding of tRNA gene and tME diversity across eukaryotes is pinned on a handful of representative model organisms ^10^. The development of next-generation sequencing technologies and bioinformatic techniques has provided a toolbox for the study of tRNA biology in previously unachievable granularity. Firstly, the increasing availability of whole genomes led to the development of tRNA identifying software tRNA-scan, with over 600 genomes on the current GtRNAdb database ^38,39^. Subsequently, the development of high-throughput gene knock-out methods allowed for experimental analysis of tRNA gene and modification enzyme essentiality, and role in some species ^9,40–42^. Further technological advancements in RNA sequencing have led to cutting-edge methods in tRNA sequencing and tRNA modification calling in eukaryotes ^43–47^.

Here, we employed whole-genome analysis of 1153 genomes from 1,050 species from the fungal subphylum Saccharomycotina (hereafter referred to as yeast) to explore eukaryotic tRNA and tME gene diversity on a previously unseen scale. We analysed tRNA gene sequence diversity across the entire subphylum and found isodecoder levels increased with genome length and tRNA gene counts. However, we observed that despite the general trend, several phylogenetic orders displayed high levels of tRNA with low sequence diversity and vice versa, suggesting contrasting evolutionary trends in tRNA evolution. Machine learning analyses found that tRNA sequence is predictive of taxonomic classification and further discovered distinct conservation of tRNA groups between taxonomic orders. Next, we annotated tME across the subphylum and found surprising patterns of loss for several tME pathways and complexes. We also found that tME repertoire was associated with genomic GC content, highlighting complex modes of tME and tRNA co-evolution in eukaryotes.

Finally, we applied cutting-edge direct tRNA sequencing technology to observe tRNA modifications across three selected Saccharomycotina species: Saccharomyces cerevisiae, Yarrowia lipolytica, and *Hanseniaspora uvarum*. We compared modified fractions of tRNA between species, focusing on sites where single modifications are known. We found that fractions of modified tRNA differed between species for certain modifications only. We then compared total modification profiles between tRNA groups across all three species and found no overall statistical difference between modification profiles. We then explored whether absence of tME was associated with absence of target nucleotides. Interestingly, we found that while differential tME repertoires did not appear to associate with differential modification profiles, tME repertoires were indicative of target nucleotide presence in tRNA sequences. These results provide a novel insight into the realm of eukaryotic tRNA modification and suggest a complex interplay between tME repertoire and tRNA sequence evolution. This work opens a range of questions regarding distinct evolutionary paths of modification between species.

The genetically diverse Saccharomycotina subphylum provides an experimental and genomic treasure trove for exploring tME repertoires, which is more representative of eukaryotic diversity. Our current understanding of tRNA biology has been dominated by a handful of species; this has led to ‘universal’ theories across all eukaryotes. This study strongly challenges what we know about tRNA in eukaryotes and brings about a new era of interspecies analyses that will change our understanding of the fundamental biology of life.

## Results

### Saccharomycotina tRNA isotype diversity and taxonomic distribution display lineage-specific patterns

The highly conserved structure of tRNA genes could suggest a lack of nucleotide diversity that would be under selection to maintain their integral function. However, studies of tRNA gene sequence diversity in a handful of eukaryotic organisms have shown tRNA to be diverse in nature, though these examples remain limited ^10,48^. Here, we have carried out a tRNA nucleotide diversity analysis by analysing over 200,000 tRNA gene sequences belonging to the Saccharomycotina subphylum (Fig. 1 a).

**Figure 1.**
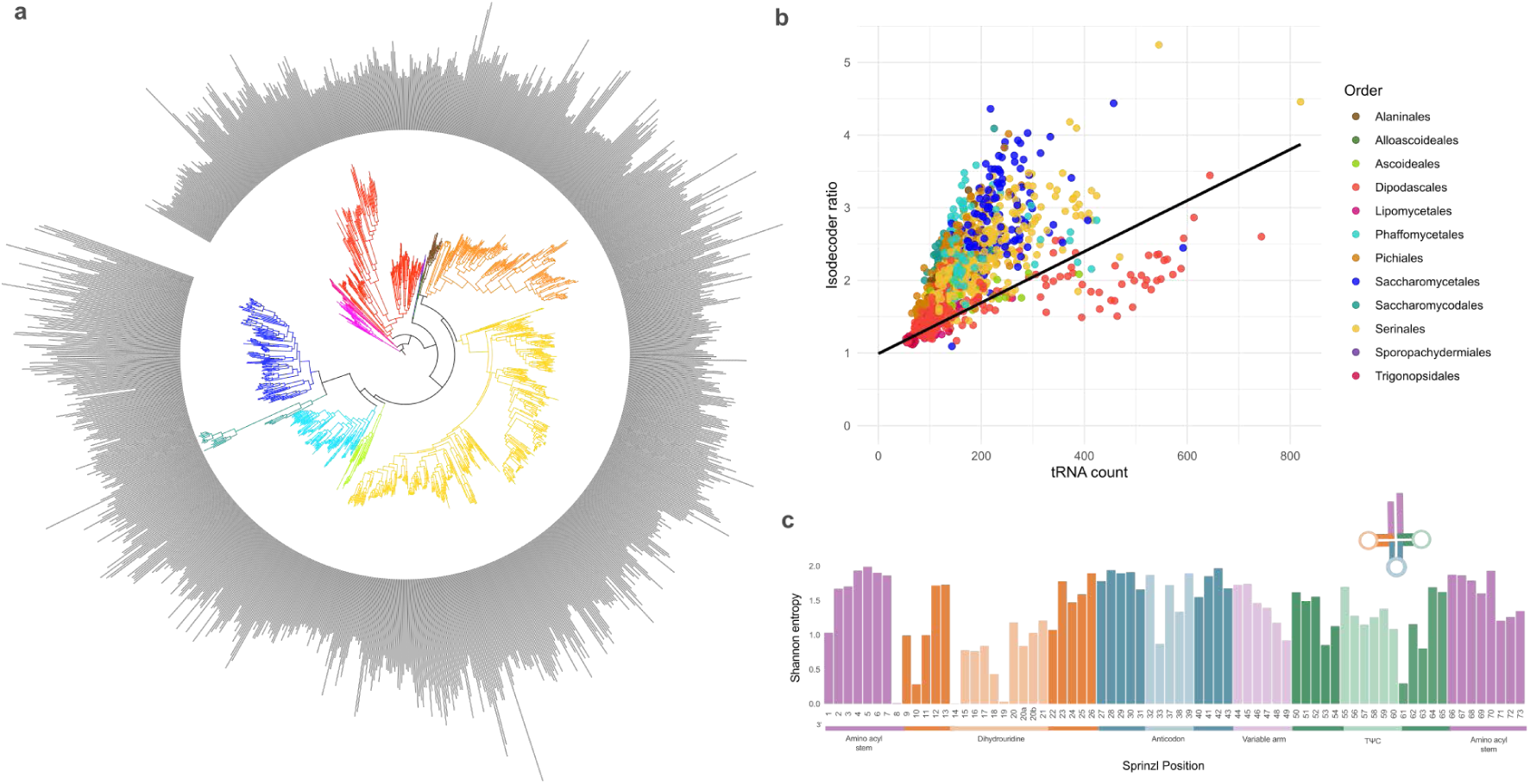
tRNA gene sequence diversity analysis. a) Phylogenetic tree of Saccharomycotina with a bar chart displaying total tRNA gene count per species. b) Scatterplot showing isodecoder ratio vs total tRNA count for each species, coloured by phylogenetic order. c) Bar chart showing Shannon entropy values at each Sprinzl position across all tRNA genes of the Saccharomycotina

We focused on unique tRNA isodecoder ratio as a measure for total tRNA gene sequence diversity within a genome. tRNA isodecoder ratio was defined as the total number of unique isodecoders per tRNA gene within a species. For example, *S. cerevisiae* has 276 tRNA genes, within these genes only 71 unique isodecoder sequences are present, this is an isodecoder ratio of 3.89. A large range of isodecoder ratios was found within the Saccharomycotina (1.09 in *Aciculoconidium aculeatum*, 5.24 in *Zygosaccharomyces bailii*), with an average ratio of 2.2. Using a Phylogenetic Generalized Least Squares (PGLS) we determined that isodecoder ratio and total tRNA gene count show a significantly positive correlation when all species are considered, and that phylogeny accounted for similarity between species (PGLS: R2 = 0.51, p = 0, λ = 0.940 Fig. 1b). Species within the Dipodascales order did not follow the overall trend and had some of the highest tRNA counts in the subphylum but did not display an increase in sequence diversity as count increased. For example, the species *Magnusiomyces magnusii* has 744 genes and a ratio of only 2.6. This result suggests that tRNA sequence diversity in the Dipodascales was subject to stabilising selection. Across the rest of the Saccharomycotina, however, the correlation between tRNA sequence diversity and tRNA gene content suggests that expansion of isodecoder ratio is facilitated by increased gene copy number and therefore genome size, as reported in previous work ^49^.

Next, we examined position-specific diversity within tRNAs as we suspected that certain tRNA positions are likely to be more conserved than others due to roles in aminoacylation and tertiary structure. We calculated Shannon entropy values for all positions across all tRNA genes in the subphylum (Fig. 1c). We found that a small number of positions show low diversity across all genes and genomes; 8, 14, and 19 which belong to the 3’ final nucleotide of the aminoacyl stem and Dihydrouridine region. Interestingly, a trend of increasing diversity can be seen leading up to the anticodon region from both the 3’ and 5’ ends, and the aminoacyl stem also shows relatively high nucleotide diversity (Fig. 1b). The conservation of positions 8 and 14 is observed across domains and is due to reported interactions between these bases during tertiary folding ^50^. High nucleotide diversity of the aminoacyl stem likely results from aminoacylation recognition selecting for specific nucleotides ^51,52^. Additionally, the helical structure of the stem is constrained, leading to the allowance of nucleotide pair diversity ^53^.

We hypothesised that tRNA nucleotide diversity may be evolutionarily conserved. To test this, a Random Forest model was then employed to test if a tRNA sequence within a given amino acid group could be classified into the correct taxonomic order based on sequence alone. We found that the model can correctly classify unseen tRNA genes into the correct order with high accuracy for most orders (Fig. 2a). tRNA belonging to the Lipomycetales was generally correctly assigned (61.7%); however, tRNA could also be assigned to the closely related Trigonopsidales and Dipodascales orders despite a high accuracy (99-100%) for tRNA belonging to these orders. This was an interesting result, as we had previously seen that species in the Dipodascales order had high tRNA gene counts, with low isodecoder diversity. This suggests that in Dipodascales, conserved bases are a defining characteristic of the order that enables classification. In Lipomycetales, the results suggest that tRNA are diverse and do not contain conserved changes.

**Figure 2.**
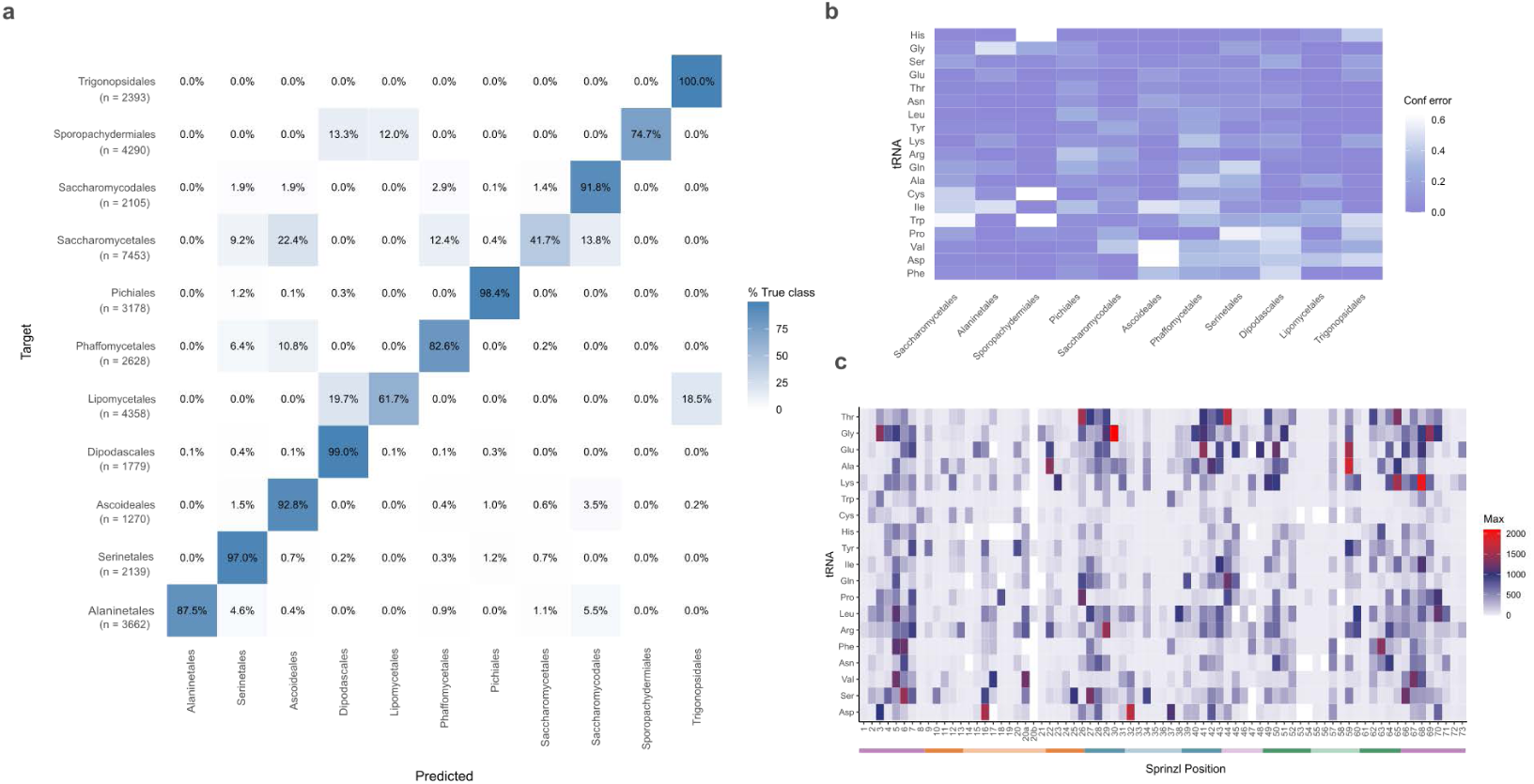
Random Forest classification analysis of aligned tRNA sequences. a) confusion matrix of Random Forest results of tRNA within isotype group classification to phylogenetic order. b) Heatmap showing confidence error of the Random Forest model within each tRNA isotype group. c) Heatmap showing feature importance in each mode using Mean Decrease Gini for each feature at each position. The maximum feature importance per amino acid per position is reported.

The data generated in our Random Forest model was then analysed further to investigate whether certain tRNA groups were predicted with more confidence than others (Fig. 2b). Our results indicated that successful classification of tRNAs was highly differential between amino acid groups and order. For example, tRNA carrying Trp, Pro, Val, Asp, and Phe amino acids are classified with less confidence in orders Phaffomycetales, Serinales, Dipodascales, Lipomycetales, and Trigonopsidales. Nevertheless, based on sequence alone, we would be able to correctly classify a novel tRNA into the correct taxonomic order for most amino acids.

Finally, we predicted that certain positions of tRNA may also drive accurate classification in the models within amino acid groups. We observed that several positions were highly informative in the model, dependent on amino acid grouping (Fig. 2c). Interestingly, the positions that were most important to the model were distributed across the entire structure. This suggests that phylogenetic signals within the tRNA gene sequence are not specific to a particular region. We observed that the areas showing a trend of importance were the aminoacyl stem and the anticodon stem. When considering the tertiary structure of a tRNA, these two regions form the arm. The Ribothymidine-Pseudouridine-Cytidine loop (T-loop) and dihydrouridine loop (D-loop) interact to form the structural elbow of a tRNA, which may explain the low importance as the tertiary structure is integral for the function of tRNA, therefore conservation of these regions is expected. The aminoacyl and anticodon stem, however, do not form structural interactions. A critical step in preprocessing of tRNA is aminoacylation and post-transcriptional modification. These enzymes interact with both the aminoacyl and anticodon stem to discern between tRNA type. The dependence of enzymes on the interaction with these regions may explain high importance across amino acid groups. Interestingly, we do see that no Sprinzl positions of Lys, Trp, Cys, His, Tyr and Ile tRNA had have high maximal importance to the model. Conversely, there is at least one Sprinzl position within each of the remaining amino acid groups that has maximal importance to the model when predicting phylogeny. Divergence of certain nucleotides between orders could be a signal of lineage-specific isodecoders, where diversity is permissive in comparison to other nucleotides.

Despite conserved function across all life, our analysis of tRNA diversity in the Saccharomycotina subphylum found high levels of sequence divergence across phylogenetic orders, providing evidence of complex tRNA-associated evolution in eukaryotes. Several critical questions are highlighted regarding fundamental tRNA biology in eukaryotes, such as why tRNA sequences are phylogenetically distinct within certain tRNA types but not others, and what evolutionary forces are driving such distinctions. Further explanations for such distinctions may also be linked to other non-canonical regulatory functions of tRNA, such as genome architecture and undiscovered regulatory roles.

### tRNA modifying enzyme repertoire size across the Saccharomycotina are diverse

To characterize the diversity of tMEs, we annotated the tME repertoires across all 1154 genomes in the Saccharomycotina subphylum. The tME repertoire was defined as a total of 83 enzymes currently known to exist in fungi, including pre-processing enzymes. We hypothesised that the diversity of tMEs would be variable across the subphylum, and that not all Saccharomycontina yeasts would carry a universal tME repertoire. Using a Hidden Markov Model (HMM), the Saccharomycotina subphylum genomes were re-annotated to capture a robust tME presence dataset. We identified a total of 11,759 tMEs across all genomes and found no instance of all 83, only 17 species were found with the maximum total of 80 enzymes known in fungi.

tME repertoire size was diverse across the subphylum, with a range of 60-80 enzymes (Fig. 3a(i)). Closely related orders Lipomycetales (75), Trigonopsidales (73), Dipodascales (73), Sporopachydermiales (75), and Alloascoideales (75) have the largest repertoires on average within the subphylum. Most drastically, the order Saccharomycodales had the lowest average (66), and the largest range of repertoires belonged to the Serinales order (60-79). The total number of tMEs is significantly diverse between phylogenetic orders (Fig. 3b) (ANOVA, df=11, F=60.69, p<0.001), and only 48 tME are conserved in over 95% of Saccharomycotina species. These findings demonstrate that a tME repertoire for any given species within the Saccharomycotina can range from 60 to 80, and that phylogenetic order is not predictive of tME repertoire size.

**Figure 3.**
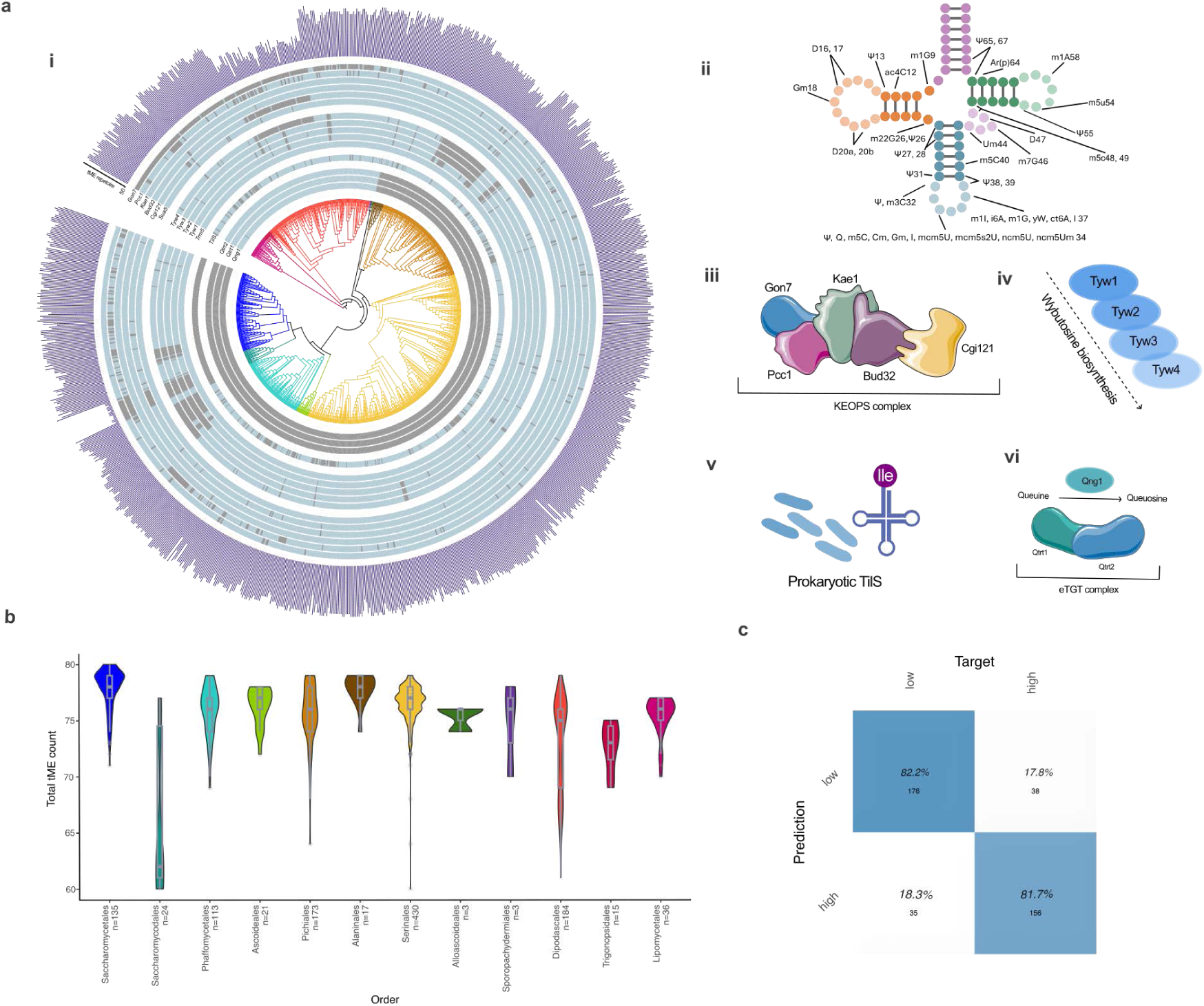
tRNA modification enzyme annotation analysis of Saccharomycotina subphylum. a) i) Presence or absence of highlighted tME across the Saccharomycotina subphylum phylogenetic tree. Bar chart plot of total tME count per species, the y-axis is truncated at 50. Components of the KEOPS complex presence or absence. Wybutosine biosynthesis pathway enzyme presence or absence. tilS presence absence. Queuosine salvage and eTGT complex components. ii) tRNA molecule annotated with modifications known to occur in fungi. iii)-vi) Illustrative figures of tME highlighted. b) Violin plot showing distribution of total tME gene counts within each order, boxplots indicate median and interquartile range. The number of species within each order is indicated on the x-axis. c) Confusion matrix of Random Forest classification of genomic GC content using tME repertoire.

Interestingly, we see a distinct pattern of absence across the Saccharomycodales order for several tME. Previous studies found that within this clade, a faster-evolving lineage occurs, and that several DNA mismatch repair genes were also absent from this lineage ^54^. We also found that the average total tME repertoire for this order is the lowest for a complete order (Fig. 3b). The loss of tME in Saccharomycodales is consistent with previously reported gene loss and likely has a huge impact on the genome. Further investigation into the tRNA biology of these fungi may provide a novel insight into the translation mechanics of cells lacking several canonical tME involved in translation maintenance.

### Gain and loss of tRNA pathways is common across the Saccharomycotina

Many tRNA modifications require protein complexes and multi-enzymatic pathways for synthesis ^22^. Therefore, tME presence and absence data were grouped by complex and enzymatic pathway to identify complete modification pathways. We focused on a set of complexes and enzymatic pathways that showed distinct patterns of loss/conservation across the subphylum (Fig. 3a(i)). First, we examined the KEOPS complex (Kinase, Endopeptidase and Other Proteins of Small size), which is made up of a putative kinase Bud32 alongside Gon7, Pcc1, Cgi121, and Kae1 (Fig. 3a(iii)) ^55^. Together with Sua5, the KEOPS complex synthesizes the N6-threonylcarbamoyladenosine (t6A) at position 37 on the anticodon loop on ANN decoding tRNA ^56^. The initial step in the biosynthesis of t6A requires Sua5, our annotation analysis found that this tME gene was highly conserved across the subphylum. The enzymes comprising the KEOPS complex displayed a complex pattern of conservation. Specifically, *bud32* and *kae1* were largely present across the subphylum. Conversely, *cgi121*, *pcc1*, and gon7 were not predicted in several species belonging to the Dipodascales, Saccharomycodales, Trignopsidales, and Lipomycetales orders. Our findings of enzyme-dependent conservation of KEOPS components align with current literature, specifically, Pcc1 and Gon7 are predicted to play a supporting role within the complex, Pcc1 is reportedly not required for catalytic activity ^57^, whereas Kae1 and Bud32 are essential for t6A reaction ^55,58^. The significance of the conservation pattern of *cgi121* is less clear, as recent findings suggest that Cgi121 is required for tRNA binding ^56^.

The second pathway examined was the wybutosine enzymatic pathway, which synthesizes a wybutosine (yW) modification derived from guanine at position 37 of Phe-tRNA. The biosynthesis requires Trm5 to catalyse the initial modification of G37, which is further modified by an enzymatic pathway comprised of enzymes Twy1-4 (Fig. 3a(vi)) ^59^. Our prediction found that the entire pathway is absent from numerous species across the subphylum spanning several orders: Saccharomycodales, Pichiales, Phaffomycetales, and Saccharomycotales (Fig. 3a(i)). We found that, in contrast, *trm5* is widely conserved. Trm5 methylates G37 and has been shown to methylate other tRNA positions ^60^, this moonlighting function of Trm5 could explain the conservation pattern observed in our dataset.

We also searched for evidence of the queuosine (Q) modification enzymes acting on the guanine at position 34 of NAU decoding tRNA (Tyr, His, Asp, and Asn) ^61^. The free base of Q, queuine, replaces the guanine via the process of transglycosylation catalysed by the heterodimer tRNA guanine transglycosylase (Qtrt1/Qtrt2(/d1)) ^62,63^. Eukaryotes must salvage queuine (Q nucleobase) from the environment (via Qng1), whereas certain bacterial species synthesise queuine ^62,63^. We find that 5 phylogenetic orders within the Saccharomycotina have Q-tME and the queuine salvage protein *qng1*: Lipomycetales, Trigonopsidales, Dipodascales, Alloascoideales, and Sporopachydermiales (Fig. 3a(i)). The ability to engage in queosine modification appears to have been lost in the Saccharomycotina approximately 311 million years ago ^64^. We also employed our HMM annotation method to find evidence of the hyper-modifiers of Q, *manQ,* and *galQ,* recently defined in humans ^65^. Our analysis did not identify hyper Q modification enzymes in the subphylum. Previous work has shown that loss of Q tMEs can result in an imbalance of translation speeds between GC and AU rich codons in eukaryotes. To test if GC content was associated with Q tME presence we carried out a phylogenetic generalized least squares analysis (PGLS) and found that presence of the Q tMEs shows no relation to GC content when phylogeny is considered (R-squared: 0.001, p-value: 0.56). However, when visualising the distribution of GC content across all orders, we observe a higher average GC content in the orders with conserved Q-tME’s (Supplementary Fig. 1). Despite not identifying a statistical relationship between Q-tME’s and GC content, we further questioned whether total tME repertoire could be indicative of GC content. To test this, a Random Forest model was employed (Fig. 3c). We find that the tME repertoire could be used to classify high and low quantiles of GC content for Saccharomycotina species with high accuracy.

### Identification of previously uncharacterized tMEs in fungi

We further speculated that tME, known to occur in mammalian species, or modifications that are not found in model Saccharomycotina species, may also be present in the subphylum. Using the HMM annotation method described and applied above, we searched for evidence of additional tMEs *trdmt1*, *trmo*, *mtaB,* and *alkbh1* in our dataset and found evidence for the presence of *alkbh1*. The orders in which we identified at least one copy of *alkbh1* were Lipomycetales, Trigonopsidales, Dipodascales, Alloascoideales, and Sporopachydermiales, following the same pattern of conservation as the Q tMEs. Alkbh1 is an RNA demethylase belonging to the human AlkB family dioxygenases and acts to demethylate the *N*^1^-methyladenosine (m^1^A) modification at canonical position 58 ^66^. An *alkbh1* homolog has previously been annotated in *Magnaporthe oryzae* ^67^ and *Schizosaccharomyces pombe* ^68^ and remains the only fungal homolog reported in literature. Interestingly, the *S. pombe* homolog reportedly does not have demethylase activity.

We then applied the HMM method and searched for the presence of 20 tME known in prokaryotes. We find novel evidence of one tME known only to exist in prokaryotes, TilS (Fig. 3a(iv)). *tilS* shows extensive loss across the Saccharomycetales and Saccharomycodales orders. Presence of *tilS* found in this analysis has not been previously reported in the literature in fungi, however it has been reported as a candidate mitochondrial enzyme in *Arabidopsis thaliana* ^69^. Previous work found no homolog of *tilS* in mammalian genomes, this was suggested to be because the AUA codon is decoded by Ile tRNA with inosine/pseudouridine at wobble positions ^70^. The same is true for the modification of Ile-tRNA in *S. cerevisiae* ^71^. Interestingly, we do not find evidence of *tilS* in the Saccharomycetales; however, it is likely that, like *A. thaliana*, *tilS* is a mitochondrial tME encoded in the nuclear genome.

The initial findings in our tME repertoire analysis present a contrast to current literature. The KEOPS complex is considered universal across eukaryotes at present, with *gon7* homologs found in fungi and humans ^72^. Our analysis finds that *gon7* is likely not present in all fungi, and that the pattern of conservation is more complex than between kingdom conservation. Additionally, the lack of multiple enzymes comprising the KEOPS complex may result in a lack of t6A modification or comparatively could provide insights of essentiality and diversity in the biosynthesis of t6A across species. These results also highlighted a strong distinction of conservation patterns between the presence *of alkbh1 and Q, and the divergence of orders from the last common ancestor between Lipomycetales, Trigonopsidales, Dipodascales, Alloscoideatales,* and Sporopachydermiales, and the rest of the subphylum. The pattern of conservation observed suggests widespread loss across the phylogeny, with the ancestor retaining the most enzymes.

### Cross comparison of post-transcriptional modification using direct RNA sequencing

We hypothesized that the genomic tRNA sequence and tME diversity we observed across the subphylum would be reflected in the *in vivo* tRNA modification profile diversity. At present, tRNA sequencing has not been carried out across a diverse set of eukaryotic species in parallel. We applied a direct RNA sequencing approach developed by Lucas et al. (2024) and further by White et al. (2025) ^47,73^ to sequence tRNA in three genetically diverse species of Saccharomycotina: Saccharomyces cerevisiae, Hanseniaspora uvarum, and *Yarrowia lipolytica* (Fig. 4a). A total of 385680 reads spanned all three species, *Saccharomyces cerevisiae* (36.5%), *Hanseniaspora uvarum* (48.8%), and *Yarrowia lipolytica* (14.7%). tRNA modifications were predicted using a method of quantifying post-alignment base call (BC) error above a background threshold.

**Figure 4.**
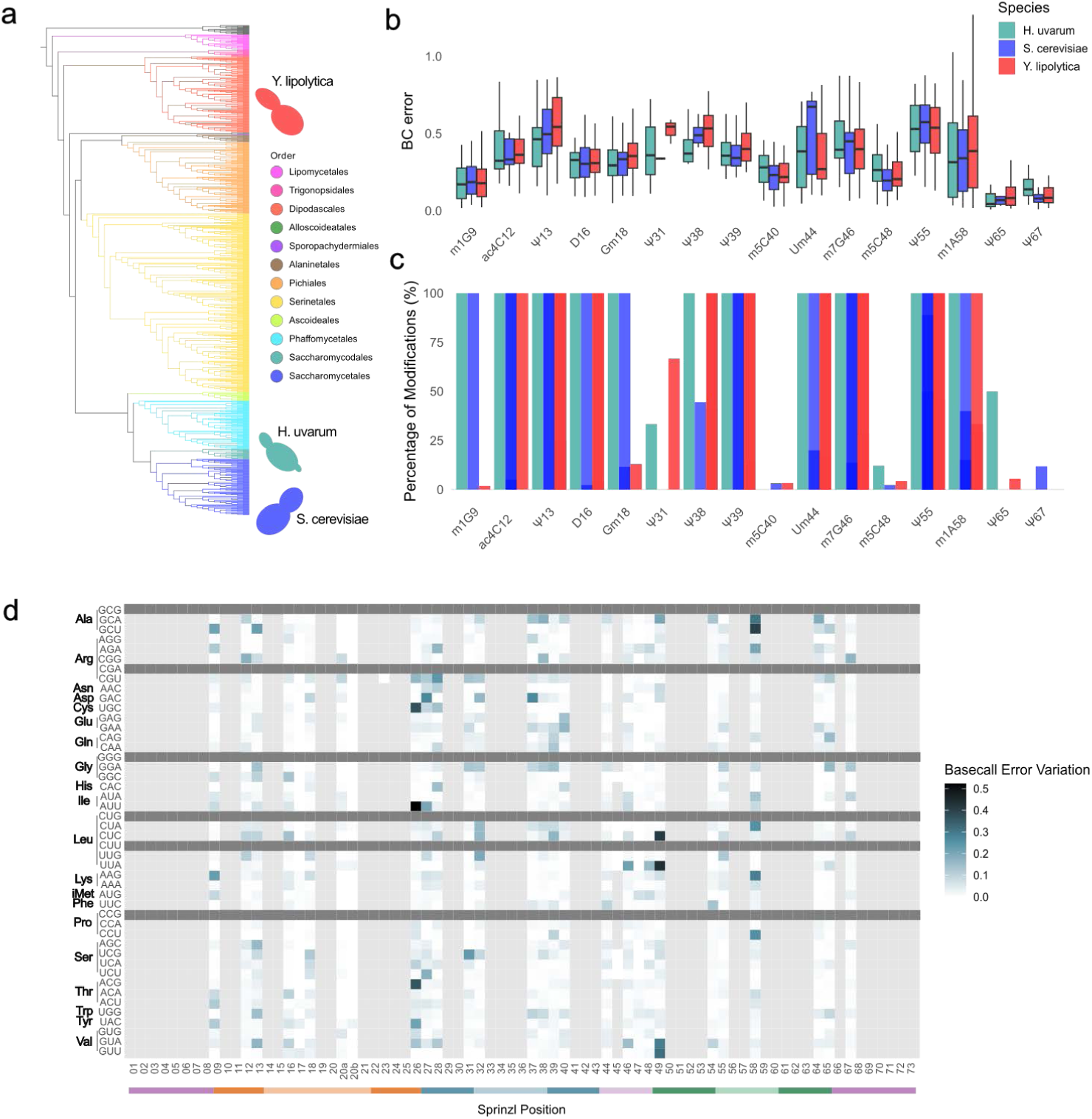
Direct tRNA sequencing of three Saccharomycotina species. a) Saccharonycotina subphylum phylogenetic tree labelled with tRNA sequenced species *Hansaniaspora uvarum*, *Yarrowia lipolytica* and *Saccharomyces cerevisiae*. b) Boxplot showing raw BC error values for single modification sites for each species. c) bar plot showing the fraction of modified tRNA at specified modification sites between species. d) heatmap showing variation of BC error within tRNA group between species.

To test whether the fraction of modified tRNA at specific sites would differ between species, we focused on modification sites that were known to harbour single modifications. Using our aligned tRNA gene dataset, we filtered for genes with the modifiable nucleotide present at the canonical position and compared BC error signals across species at these sites (Fig. 4b). First, we compared the BC error signal for each modification across species to explore whether this differed between the species. We observed a range of BC error values for each modified site for each species. Several modified sites displayed highly similar average BC error signals across species; m1G9, D16, D20a, Ψ55. Several modifications observed had two species with similar average BC error values; Ψ13, Ψ38, m5C40, D47, m1A58, Ψ65, and Ψ67. While the modifications ac4C12, Um44, m7G46, and m5c48 displayed variable average signals between species, an ANOVA of BC error values between species for each modification found that only m5c48 showed a significant difference (F-value = 3.98, p = 0.0208). Analysis of BC error has been shown to pick up distinct signals between modifications at the same site; smaller changes, as seen here, could also reflect chemical variations of modifications termed derivatives, or intermediate modifications ^36^. Known chemically distinct derivatives include yW and Q ^74,75^, changes to the composition of modifications due to environmental stress has also been described ^76^. Whereas several modification signals are similar between species, the differential signal for certain modifications has not previously been reported in eukaryotes. This analysis highlights the importance of studying tRNA biology from an interspecies species perspective to support modification calling models.

Next, we tested whether the fraction of modified tRNA would differ between species. We calculated percentage of modified tRNA between species (Fig. 4c). Our analysis found that while several tRNA were consistently modified across species, some tRNA showed inter-species variation. A Fisher’s exact test found no significant difference between species for each modification. These results suggest that certain modifications are required universally for tRNA function, whereas others show relative importance to environmental niches or cellular states. Modifications consistently observed may be more efficiently synthesised, whereas more variable modifications may require specific substrates, not readily available in the cell, for example. We also considered that the culture of these three species was carried out in parallel under the same conditions, as these species are genetically diverse, we may also be detecting signals of differential responses to the growth conditions.

Additionally, we hypothesised that the presence of modifications across all modified sites would vary between tRNA types. To test this, we calculated the interspecies variation of BC error at each modified canonical position within each tRNA anticodon group (see Fig4D). Our analysis found evidence for differential modification profiles between species for all comparable tRNA groups. As we found that species within the Saccharomycedales order had the smallest tME repertoires, we employed a Generalised Linear Model to test for differences in modification signal across all positions in *H. uvarum* (Saccharomycedales) compared to *S. cerevisiae* and *Y. lipolytica*. The GLM found no significant effect of species on the modification signals between the species (*S. cerevisiae* coef: -0.013, *Y. lipolytica* coef: 0.005). We then carried out an ANOVA at all positions within anticodon groups where enough tRNA genes were present. The ANOVA found that 8 anticodon groups of tRNA displayed significantly different (p<0.05) BC error between species at various positions (Supplementary table 10). For example, GAC tRNAs had 19 positions that differed significantly, whereas AGC tRNA had a single position that varied significantly in modification signals between species.

Finally, we directly compared tME absence with modification signal *H. uvarum* at positions where only a single modification is known to occur. We predicted that the tME *trm3* (Gm18) was absent in HMM analysis, comparatively we found a modification signal (BC error > 0.5) at this position (see Fig4C). We also predicted that the biosynthesis pathway for Wy37 was absent from *H. uvarum*, interestingly we found that there are no tRNA genes with a G at position 37. We suspected that the detection of Gm18 in *H. uvarum* despite predicted absence of the *trm3* gene may be due to sequence divergence resulting in an inability of the model to predict an ortholog.

### Multi-model analysis links tME repertoires with tRNA gene sequence and genomic GC content

The specificity of tMEs can vary between a canonical tRNA position or nucleotide base, in most cases, both base and position dictate tME specificity. The annotation of tMEs and tRNA gene sequence across the Saccharomycotina subphylum found a range of tME repertoires and sequence diversity. As our results above highlighted that loss of tME correlated with a loss of the target nucleotide. Based on tME specificity, we hypothesized that tME repertoire may impact tRNA gene sequence across the subphylum. For example, would the lack of a tME targeting an A base at position 37 in any given species mean that tRNA would be less likely to have A37 in each tRNA? A mixed-effects logistic regression was employed to model the effects of tME absence on tRNA nucleotides at specific canonical tRNA positions.

The wybutosine modification at position G37 on Phe-tRNA requires a biosynthetic pathway (*tyw1, tyw2, tyw3, tyw4*). A mixed-effects logistic regression finds that a Phe-tRNA belonging to a species lacking the entire biosynthetic pathway are significantly more likely to lack a G at position 37 (intercept estimate=9.8994, z=<2e-16 ***). This result correlates with our initial finding that in *H. uvarum*, Phe tRNA do not have G37 and no wybutosine tMEs were annotated. We find a similar result when examining an enzyme complex. The tA37 modification requires the KEOPS complex and modifies ANN decoding tRNA. We find that for ANN decoding tRNA, the likelihood that the tRNA will have an adenine at position 37 (A37) increases significantly when all KEOPS complex genes are present (intercept estimate=-2.28261, z=< 2e-16 ***). We further find that the presence of enzyme *gon7* significantly increases the likelihood that an ANN tRNA will have A37 (z=< 2e-16 ***).

Our multi-modal analysis suggests that tME repertoire is intrinsically linked to tRNA gene sequence evolution and genomic GC content. The tRNA processing pathway is highly regulated; for example, hypomodified or tRNA lacking modifications are readily degraded ^77^. This would suggest that loss of tMEs could have adverse effects, however if coupled with a loss of target tRNA this loss could be less disadvantageous. Additionally, tRNA modifications require various enzymatic substrates that will inevitably be limited by environmental niche. The Q-tME are an example of a nutrient driven modification, as the acquisition of the precursor, queuine, is obtained from bacteria, as they can readily synthesise queuine de novo. ^78^. The range of environmental niches inhabited by Saccharomycotina are vast and numerous and could largely explain the co-evolution and diversity of tRNA gene sequence and tME’s suggested by our findings.

## Conclusion

Until now, the diversity of tMEs in fungi, and more broadly eukaryotes, has remained largely unknown. Additionally, the current literature reflects previous unavailability of large and diverse genome sets for such analyses. Direct, cross-species comparisons of tRNA modification profiles remain rare, particularly outside bacteria and archaea ^79,80^. Here, we provide one of the first such comparisons in eukaryotes, contributing to a growing but still limited body of work in this emerging area of tRNA biology. The annotation method provides a robust high-throughput prediction of presence and absence of tME across 1154 genomes within an entire subphylum, and loss of entire pathways observed supports our gene annotation results suggesting evolutionary loss. We have also shown that modification of tRNA can be distinct between species. tMEs are ancient and evolved alongside tRNA in the early beginnings of life. Therefore, it is imperative to consider not only the genetic diversity within the Saccharomycotina but also the diversity of environmental niches that these yeast species have inhabited since diversifying approximately 400 million years ago ^64^. We have found evidence of evolutionary losses of these ancient enzymes across a highly diverse fungal subphylum, and evidence to show co-evolution of tRNA gene sequence. This work has implications in the field of fundamental tRNA biology for all life on earth. As novel tRNA modifications continue to be discovered ^79,80^, known modifications are being identified at previously unrecognized positions ^81^, and tME specificities are shown to vary between species ^82^. Together, these findings point to a new era in tRNA biology, in which phylogenetically distinct and dynamically evolving modification landscapes add an unexpected layer of regulatory and evolutionary complexity.

## Materials and methods

### tRNA alignments and Sprinzl positions calculation

The tRNA sequences were previously predicted from 1,154 yeast genome assemblies using tRNAscan-SE 2.0.9 using the standard eukaryotic parameters ^64^. To align the tRNA sequences within each isodecoder family, we first removed all introns and positions without a base call (N). All tRNA sequences in the same isodecoder family were joined together with the reference sequences. Reference sequences obtained from tRNAviz ^83^. Only Saccharomycotina references were used when possible. Four isodecoders (Ala-GGC, Asp-ATC, Gly-ACC, and Pro-GGG) did not have a Saccharomycotina reference, and any available sequences were used. The tRNA sequences were then aligned using infernal 1.1.5 ^84^. The Stockholm-formatted alignments were then converted to a fasta file using the stockholm2fasta.pl script (https://gist.github.com/mkuhn/553217). The fasta files were then converted to a sequence matrix for further analysis. To properly assign each tRNA position to a Sprinzl position ^85^, we created a custom script. Briefly, each aligned sequence was compared to the Sprinzl positions reported in tRNAviz. The most common discrepancy seen in position assignments was between 20a and 20b. Positions without Sprinzl assignments were subsequently removed from the sequences. Isodecoder analysis ratio analysis was carried out in R 4.3.3, a single *Candida orthopsilosis* genome was removed in this analysis due to naming inconsistencies.

### Machine learning classification

We hypothesized that tRNA sequences could be classified into the correct taxonomic order based on sequence data alone. One analysis was run for each set of isoacceptors that decode a single amino acid. The two smallest orders, Alloascoideales (3 species) and Sporopachydermiales (3 species), were removed to avoid generating extremely unbalanced datasets. All analyses were run using R 4.3.3 using the packages caret (v6.0.94 ^86^), randomForest (v4.7.1.1 ^87^) and iml (v0.11.4 ^88^). To build the random forest models, we used 70% of the data for training and 30% for testing. We then conducted 5-fold cross-validation repeated 10 times for 50 total resamples. Sampling was conducted using synthetic minority oversampling (SMOTE ^89^) to address class imbalance in the model. Features with zero variance were removed from the analysis. The models were then trained, and a final model was selected using accuracy. We then interrogated the models using AUC-ROC curves per fold and in the top model. We also calculated the feature importance in each mode using Mean Decrease Gini for each feature at each position. The maximum feature importance per amino acid per position was reported. To assess model performance in the testing dataset, we generated confusion matrices and calculated accuracy in each taxonomic order.

### tRNA modification enzyme annotation

To re-annotate tMEs, the functional annotations from the KEGG database ^90^ in the Saccharomycotina ^64^ were filtered for tME KEGG profiles. Gene sequences within each tME KEGG group were aligned using default settings in MAFFT (v7.525) ^91^. Alignments were then used to build a Hidden Markov Model profile and search against annotated genomes using HMMER (v3.4) ^92^. The output for each tME KEGG was then manually interrogated for an individual e-value cut-off. Genes above the e-value cut-off were then assigned as present, and genes below were assigned as absent. Presence and absence were plotted onto the phylogenetic tree using iTOL ^93^

### Multi-model analysis of annotation data and statistical testing

The Phylogenetic Least Squares model was run using R 4.3.3 using the pgls function in the caper package (v1.0.4) and used published GC content data of species and phylogenetic distances ^49,64,94,95^. Mixed effects logistic regression was done using aligned tRNA gene sequences associated with each species and tME annotation data using R 4.3.3 with the lme4 (v1.1-37) package ^96^. All statistical analyses were done in R 4.3.3.

### Strains and culture conditions

The *Saccharomyces cerevisiae* strain (CBS1171), *Hanseniaspora uvarum* (CBS 314) and *Yarrowia lipolytica* (CBS 6124) were obtained from CBS strain collection. To culture from -80 stocks, a loop of each strain was cultured overnight at 30°C with rotation in liquid YPD medium. To isolate RNA, cultures were then inoculated into liquid YPD at 0.001OD and grown at 30°C overnight with rotation. Cultures were centrifuged and total RNA was isolated using TRIzol**^®^** extraction.

### Direct tRNAseq library preparation

Deacylation, tRNA isolation and tRNAseq library preparation was carried out following an minorly adapted approach described by White et al. (2024) ^43^. 3-10 μg of total RNA was deacylated in 25 μg of nuclease free water was added to 25 μL of 100 mM Tris-HCl pH9 for 30 minutes at 30°C. Small RNAs were isolated using the Zymo RNA Clean and Concentrator-5 kit (Zymo Research, R1016) following the manufacturer’s instructions for isolating small RNA (17-200 nt). Splint adapters (5’ RNA splint adapter: /5/rCrCrUrArArGrArGrCrArArGrArArGrArArGrCrCrUrGrGrN, 3’ splint RNA:DNA adapter: /5Phos/rGrGrCrUrUrCrUrUrCrUrUrGrCrUrCrUrUrArGrGrArArArArArArArArArAAAA) were annealed for 15 seconds at 75°C at 10 μM in 100 μL of 10 mM Tris-HCl pH 7.5, 50 mM NaCl and 1 μL RNase inhibitor, followed by a cooling step to 25°C at 0.1°C/second. To ligate the annealed adapters, 300 ng of pure deacylated RNA was added to 1 μL 10x T4 RNA ligase 2 buffer, 4 μL 20% (w/v) PEG 8000, 2 μL 1 x T4 RNA 2 ligase, 1 μL RNase inhibitor and 1 μL of pre-annealed adapters was incubated at room temperature (RT) for 1 hour. To clean up the ligated tRNA, 1.8X volume of tRNA purification SPRI beads (BioDynami #40054) was added and incubated following the manufacturer’s protocol and eluted in 9 μL of nuclease-free water. The eluted small RNA was then ligated to the RTA adapter (Oxford Nanopore Technologies) using the adapted protocol developed by White et al. (2024) ^43^. At this stage, the samples were normalized and pooled. Then, ONT SQK-RNA004 kit instructions were followed to ligate the RLA adapter (RNA004) and clean up the library. Sequencing of the tRNA libraries was run on the PromethION P2 Solo instrument connected to a Linux workstation running MinKNOW (v24.11). The libraries were loaded onto SQK-RNA004 flow cells and sequenced using default settings ^47^.

### Basecalling and tRNA alignment

As small size RNA was filtered out using custom settings, basecalling of total raw POD5 files was re-run using Dorado (v0.8.x). Libraries were basecalled using the “super high accuracy” (rna004_130_bps_sup@v5.1.0) model, and --min-qscore 7 to quality filter reads. To make an alignment reference index of tRNA gene sequences, non-unique genes were compressed to make a unique isodecoder reference using the respective genomes. Alignment was carried out using BWA-MEM (v0.7.16) using parameters; bwa mem -C -W 13 -k 6 -T 20 -x ont2d. Reads were aligned to *S. cerevisiae* S288C, *H. uvarum* NRRL Y-1614, and *Y. lipolytica* CLIB 122.

### Modification calling

Modification calling was carried out using basecall error statistics. To get basecalling error statistics, the Python script get_bc_error_freqs.py developed by White et al. ^43^ was employed; see <u>github.com/rnabioco/tRNA004/</u>. A cut-off threshold of BC error 0.5 was used to filter for positive modification signals. Basecall error consensus sequences were aligned to the canonical Sprinzl positions using the tRNA alignment method described above. Basecall errors above the threshold at modified Sprinzl positions were deemed positive modification signals.

## Acknowledgements

The authors thank the members of the LaBella Lab and the Y1000 Project (http://y1000plus.org) team.

## Data Availability

The data underlying this article are available in a Figshare repository that will be made available upon publication of the article.

## Supplementary index

**Supplementary Fig 1.** Violin plot showing the distribution of GC content for each species grouped by phylogenetic order.

**Supplementary table 1.** Isodecoder count, tRNA count, isodecoder ratio and GC content for 1153 Saccharomycotina species genomes.

**Supplementary table 2.** Shannon entropy values across all species for each Sprinzl position.

**Supplementary table 3.** Random Forest confidence error results for classification of any given tRNA from an isotype group into phylogenetic clade (3.1). Random Forest error analysis results (3.2). Mean Decrease Gini results from Random Forest analysis of tRNA isotype classification (3.3).

**Supplementary table 4.** Presence/absence results of HMM annotation analysis for all tMEs across all Saccharomycotina species. Total modification count is shown.

**Supplementary table 5.** tRNA modification enzymes with KEGG ID and associated modification.

**Supplementary table 6.** Species GC content quantile data.

**Supplementary table 7.** Modification fraction analysis across *S. cerevisiae, H. uvarum, Y. lipolytica*.

**Supplementary table 8.** Fisher’s exact results comparing tRNA modification fractions between species.

**Supplementary table 9.** Variation analysis results for all Sprinzl positions between species and within isotype groups.

**Supplementary table 10.** ANOVA results of comparison of BC error variation between species within codon group.

