## Supplemental Figure 1 for "Hidden Diversity in Yeast tRNAs: Comparative Genomics and Modification Mapping in a Eukaryotic Subphylum"

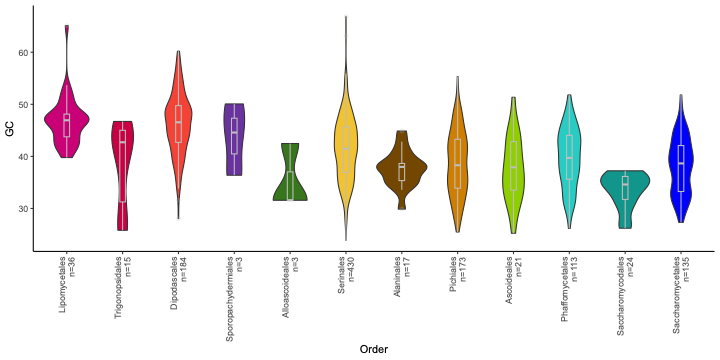

**Supplementary Figure 1. Violin plot showing the distribution of GC content for each species grouped by phylogenetic order.**
